# Caterpillars lack a resident gut microbiome

**DOI:** 10.1101/132522

**Authors:** Tobin J. Hammer, Daniel H. Janzen, Winnifred Hallwachs, Samuel L. Jaffe, Noah Fierer

## Abstract

Many animals are inhabited by microbial symbionts that influence their hosts’ development, physiology, ecological interactions, and evolutionary diversification. However, firm evidence for the existence and functional importance of resident microbiomes in larval Lepidoptera (caterpillars) is lacking, despite the fact that these insects are enormously diverse, major agricultural pests, and dominant herbivores in many ecosystems. Using 16S rRNA gene sequencing and quantitative PCR, we characterized the gut microbiomes of wild leaf-feeding caterpillars in the United States and Costa Rica, representing 124 species from 16 families. Compared with other insects and vertebrates assayed using the same methods, the microbes we detected in caterpillar guts were unusually low-density and highly variable among individuals. Furthermore, the abundance and composition of leaf-associated microbes were reflected in the feces of caterpillars consuming the same plants. Thus, microbes ingested with food are present (though possibly dead or dormant) in the caterpillar gut, but host-specific, resident symbionts are largely absent. To test whether transient microbes might still contribute to feeding and development, we conducted an experiment on field-collected caterpillars of the model species *Manduca sexta*. Antibiotic suppression of gut bacterial activity did not significantly affect caterpillar weight gain, development, or survival. The high pH, simple gut structure, and fast transit times that typify caterpillar digestive physiology may prevent microbial colonization. Moreover, host-encoded digestive and detoxification mechanisms likely render microbes unnecessary for caterpillar herbivory. Caterpillars illustrate the potential ecological and evolutionary benefits of independence from symbionts, a lifestyle which may be widespread among animals.

## Introduction

Many animals are colonized by microbial symbionts that have beneficial and fundamentally important impacts on host biology. Microbes can regulate animal development, immunity and metabolism, mediate ecological interactions, and facilitate the evolutionary origin and diversification of animal clades (1–7). These integral host-microbe relationships have led to the notion that all animals can be conceptualized as “holobionts” (8–10), superorganism-like entities composed of the host plus its microbiome—defined here as the entire assemblage of commensal, pathogenic, and mutualistic microorganisms (11). Furthermore, the recent proliferation of microbiome surveys supports a widely held assumption that microbial symbioses are universal across animals (12, 13).

The Lepidoptera (butterflies, moths, and their caterpillar larvae), despite being key components of most terrestrial foodwebs and extraordinarily diverse (14), are one group in which the role of microbes remains ambiguous. Here we focus on caterpillars, which are the main—and in some Lepidoptera, the exclusive—feeding stage, and which have long been intensively studied in many fields (15). The vast majority of caterpillars are herbivores, and some insect herbivores rely on microbes to supplement missing nutrients, neutralize toxins, or digest plant cell walls (16, 17). However, considering caterpillars’ simple gut morphology and rapid digestive throughput, it has been speculated that microbes may be unable to persist in the caterpillar gut and do not contribute to digestion (18, 19). Indeed, microscopy-based studies report no, or minimal, microbial growth within the caterpillar gut (20–22).

DNA- and culture-based investigations of caterpillar gut microbiomes have produced mixed findings, with conflicting implications for microbial involvement in caterpillar biology. Some studies report a highly abundant and consistent bacterial community (23–25), characteristics that may indicate a functional association with the host. Others report high intraspecific variability in composition, and similarity between diet- and gut-associated microbes (26–29). Inconsistencies may arise from methodological factors such as contamination of low-biomass samples (30), starvation prior to sampling, sequencing of extracellular DNA (31), and the use of laboratory-raised insects or artificial diets (27, 32, 33). Regardless of these potential biases, when studying gut microbiomes, it is often difficult to distinguish between dead or dormant passengers (“transients” (34)) and persistent, living populations (“residents” (34) or “symbionts” *sensu* (35)). Further, microbes in the latter category may be parasitic or pathogenic, as well as beneficial. While microbes were known to cause disease in caterpillars as early as Louis Pasteur’s experiments on silkworms (36), their potential importance as mutualists remains unclear.

Do caterpillars depend on gut microbes for feeding and development? To answer this question, we surveyed microbiomes of a taxonomically and geographically broad array of wild caterpillars and conducted a field-based experiment on the model species *Manduca sexta* (Sphingidae). Our analyses are focused on the digestive tract, the most likely habitat for microbial colonization, as abundant microbes have not been observed elsewhere in the caterpillar body (32, 37). First, we characterized gut microbial abundance and composition across 124 species of actively feeding caterpillars in Costa Rica and the United States. We applied the same methods to 24 additional insect, bird, and mammal species that we expected to have functional microbiomes, to assess the reliability of our protocol and to contextualize our findings. Second, we experimentally tested whether gut bacteria impact the growth and survival of *M. sexta*. Our findings question the generality of animal-microbe symbioses, and may inform a multitude of research programs based on caterpillar herbivory in both natural and managed ecosystems (e.g., (38–41)).

## Results

### Survey of caterpillar gut microbiomes

Using quantitative PCR and sequencing of the 16S rRNA gene, we found that wild caterpillars representing a broad diversity of Lepidoptera had gut bacterial densities multiple orders of magnitude lower than the whole-body microbiomes of other insects and vertebrate feces measured using identical methods (*p* < 0.0001, Fig. 1A, Table S1). Some animals host symbiotic fungi (42), but fungal biomass was also lower in caterpillar guts relative to other insects and vertebrates (median 6.1 × 10^2^ vs. 9.5 × 10^4^ rRNA gene copies per gram, *p* < 0.0001). As another line of evidence of low microbial biomass, sequence libraries from caterpillar fecal samples were dominated by plant DNA. Though there was extensive variability, typically more than 80% of 16S rRNA gene sequences in caterpillar feces were from plant chloroplasts or mitochondria, versus 0.1% for other herbivores or omnivores with plant-rich diets (*p* < 0.0001, Fig. 1B). In a subset of caterpillars from which we sampled whole, homogenized midgut and hindgut tissue, plant DNA represented an even higher proportion of sequences in guts than in feces (Fig. S1A). This pattern is more likely a function of plant DNA degradation during intestinal transit than of bacterial proliferation, as bacterial density remained similar or decreased slightly from midgut to feces, depending on the caterpillar species (Fig. S1B).

**Figure 1.**
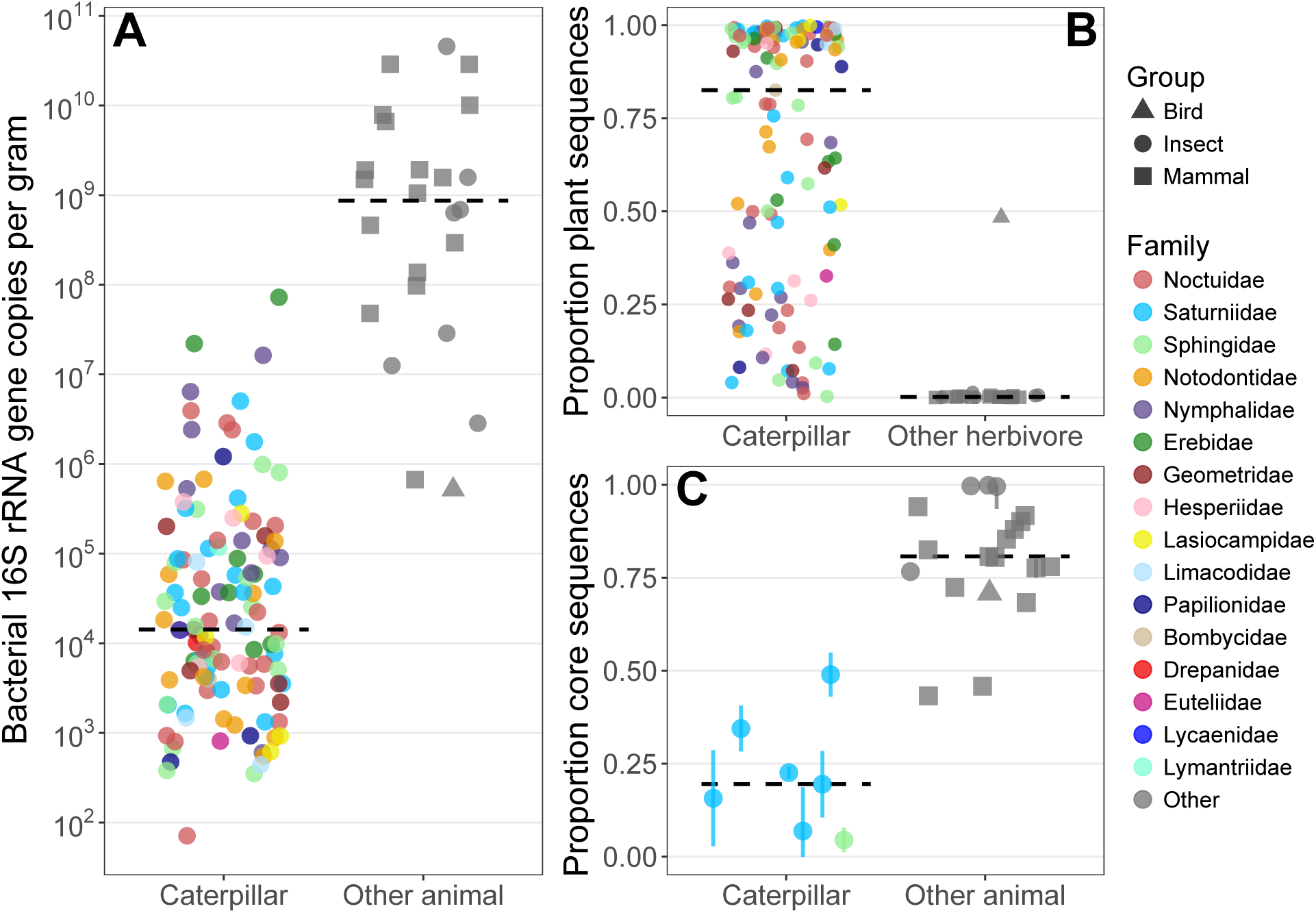
Comparisons of bacterial densities, proportions of plant DNA in sequence libraries, and intraspecific variability between caterpillars and other animals expected to host functional microbiomes. Medians are indicated by black dashed lines, and points are horizontally jittered. Data for each species are listed in Table S1. One caterpillar species yielding <100 total sequences was excluded. A) The density of bacterial 16S rRNA gene copies in caterpillar feces versus fecal (vertebrates) or whole-body homogenate (other insect) samples of other animals (N=121 caterpillar species, 24 other species). Two caterpillar species with lower amplification than DNA extraction blanks are not shown. For species with multiple replicates, the median is plotted. B) The proportion of sequence libraries assigned to plant chloroplast or mitochondrial 16S rRNA (N=123 caterpillars, 21 other herbivores). C) The proportion of sequences belonging to core phylotypes, defined as those present in all conspecific individuals. Included are species with at least three replicates with >100 bacterial sequences each (N=7 caterpillars, 19 other animals). For species with more than three, points show the median core size across all combinations of three individuals, and error bars show the interquartile range.

Caterpillar gut bacterial assemblages also exhibited a high degree of intraspecific variability, as shown by higher beta diversity within caterpillar species relative to other insects and vertebrates (*p* = 0.0002). Such variability could indicate that the microbes found in caterpillar guts are generally transient, as animals with functionally important, resident microbiomes tend to host a high abundance of microbial taxa shared among conspecific individuals (e.g., (43–45)). In agreement with this expectation, within most species of the other animals analyzed here, microbiomes were largely made up of a common set of bacterial phylotypes. For example, >99% of sequences in any one honeybee belonged to phylotypes found in all honeybees included in the analysis. In contrast, even when raised on the same species of food plant under identical conditions, caterpillars had a much lower proportion of their gut bacterial assemblage belonging to core phylotypes (median 19.5%, *p* < 0.0001; Fig. 1C). In *Schausiella santarosensis*, which among caterpillars had the highest median core size of ∼50%, four of its six core phylotypes belong to *Methylobacterium*, a typical inhabitant of leaf surfaces (46). This observation hints that many of the core phylotypes which were found in caterpillars may be transient, food-derived microbes.

In addition to low total abundance and high inter-individual variability, caterpillar gut bacterial assemblages are dominated by leaf-associated taxa, further suggesting that resident, host-specific symbionts are sparse or absent. The bacterial phylotypes present in the feces of at least half of the sampled caterpillar individuals are *Staphylococcus*, *Escherichia, Methylobacterium, Klebsiella*/*Enterobacter*, *Enterococcus*, and *Sphingomonas* (Table S2). In Colorado and Costa Rica, we sampled leaf-associated bacteria from the same plant individuals consumed by the sampled caterpillars to examine whether leaves are a potential source of these taxa. Of the aforementioned phylotypes, all but *Staphylococcus*—a potential caterpillar pathogen (47) or, like *Corynebacterium*, a transient from human skin (48)—are also among the ten most common phylotypes found in leaf samples (Table S2). Across caterpillar individuals, a median 89.6% (interquartile range: 80.2–99.0%) of fecal bacterial sequences belonged to phylotypes detected on leaves. However, bacterial assemblages were not identical between leaves and caterpillar feces (*p* = 0.001). Besides the potential growth of parasites and/or mutualists in the gut, this difference could arise from digestion filtering out subsets of the leaf bacterial community.

Transient input of leaf-associated microbes could explain the substantial variation we observed in caterpillar gut bacterial loads (Fig. 1A). Leaf bacterial densities were highly variable within (tomato) and between (milkweed, eggplant, tomato) plant species, and this variation was reflected in the feces of monarch (*Danaus plexxipus*) and *M. sexta* caterpillars feeding on them (R^2^ = 0.24, *p* = 0.03; Fig. 2A). Furthermore, bacterial densities dropped by a median of 214-fold from leaves to feces, suggesting that any potential bacterial growth within the gut is relatively minor (Fig. 2A). The extent of this reduction varied widely (from 5 to 8400-fold, Fig. 2A), possibly because of inter-individual or interspecific differences in physiological traits that may eliminate leaf microbes, such as gut pH. As with patterns in total abundance, variation in bacterial taxonomic composition among leaves and caterpillar feces was correlated (Mantel *r* = 0.28, *p* = 0.001; Fig. 2B). In other words, caterpillars consuming leaves with more distinct bacterial assemblages produce more distinct bacterial assemblages in their feces, as would be expected from a digestive system in which microbes are diet-derived and only transiently present. Moreover, this process could explain a potential relationship between host relatedness and microbiome structure, a pattern sometimes interpreted to indicate functional host-symbiont interactions (49). Specifically, although confamilial caterpillars in Costa Rica had marginally more similar gut bacterial assemblages than did caterpillars in different families (*p* = 0.053), they had also been feeding on plants with especially similar leaf microbiomes (*p* = 0.005).

**Figure 2.**
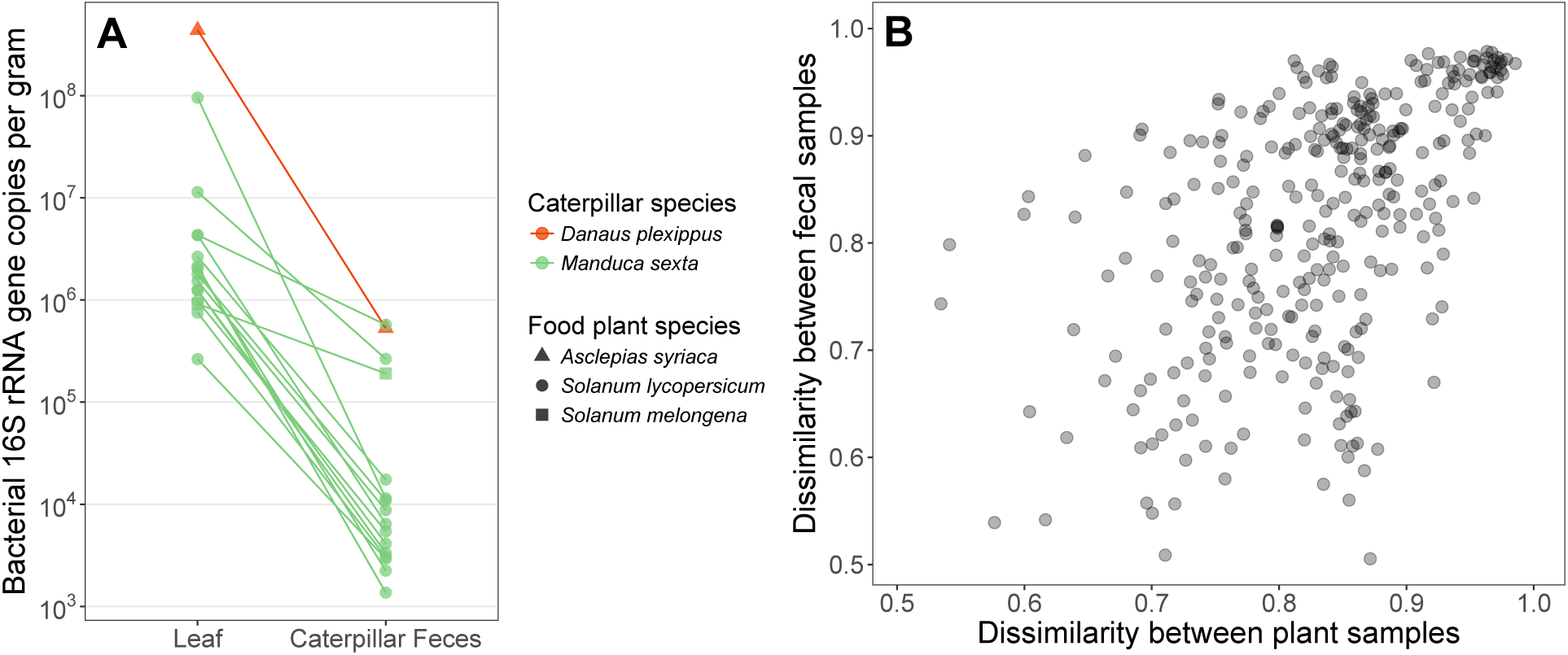
The abundance and composition of bacteria present in caterpillar fecal samples, as compared with paired diet (leaf) samples. A) The density of bacterial 16S rRNA gene copies in ground leaves versus feces, for 16 individuals collected in Colorado. Parallel lines indicate the association observed between plant and fecal bacterial abundances across paired samples. B) The correlation between beta diversity (Bray-Curtis dissimilarity metric) across caterpillar fecal samples collected in Costa Rica, and their paired leaf surface samples (N=24 caterpillar species, 19 plant species; 26 individuals each). Here only samples with >2,000 sequences are shown, to facilitate visualization.

### Test of microbiome function in Manduca sexta

Supporting our claim that caterpillars lack resident, functional gut microbiomes, we show experimentally that the growth and survival of field-collected *Manduca sexta* caterpillars are not dependent on gut bacterial activity. As measured by qPCR, wild *M. sexta* contain ∼61,000-fold lower bacterial loads than expected from allometric scaling relationships based on animals with resident microbiomes ((50), Fig. S2). Feeding *M. sexta* antibiotics reduced this already low number of gut bacteria by 14- to 365-fold (range of medians across dosages), as measured using culture-dependent methods (R^2^ = 0.13, *p* = 0.003, Fig. S3A). These colony counts were positively correlated with the number of 16S rRNA gene copies (*r* = 0.38, *p* = 0.003; Fig. S3B). Suppression of viable bacteria had no effect on pupal weight (antibiotics: *p* = 0.45; sex: *p* = 0.014; interaction: *p* = 0.70; Fig. 3), which is correlated with fecundity in insects (51), nor on development time (antibiotics: *p* = 0.19; sex: *p* = 0.023; interaction: *p* = 0.63; Fig. S4A). Likewise, antibiotic treatment did not affect survival from larval hatching to adult emergence (*p* = 0.19, Fig. S4B), nor generally impact total feces production, which is an integrated measure of leaf consumption and assimilation efficiency (antibiotics: *p* = 0.07; sex: *p* = 0.002; interaction: *p* = 0.048). As expected with *M. sexta* (52) we found clear sexual size dimorphism, suggesting our experimental design had sufficient power to detect biologically meaningful differences. Given that antibiotics reduced fecal bacteria to a variable extent within and among treatments (Fig. S3A), we repeated the aforementioned analyses using gut bacterial abundance as the predictor variable. In all cases there was no significant relationship with host performance (*p* > 0.1), further indicating that reducing or eliminating gut bacteria from caterpillars does not negatively impact *M. sexta* fitness.

**Figure 3.**
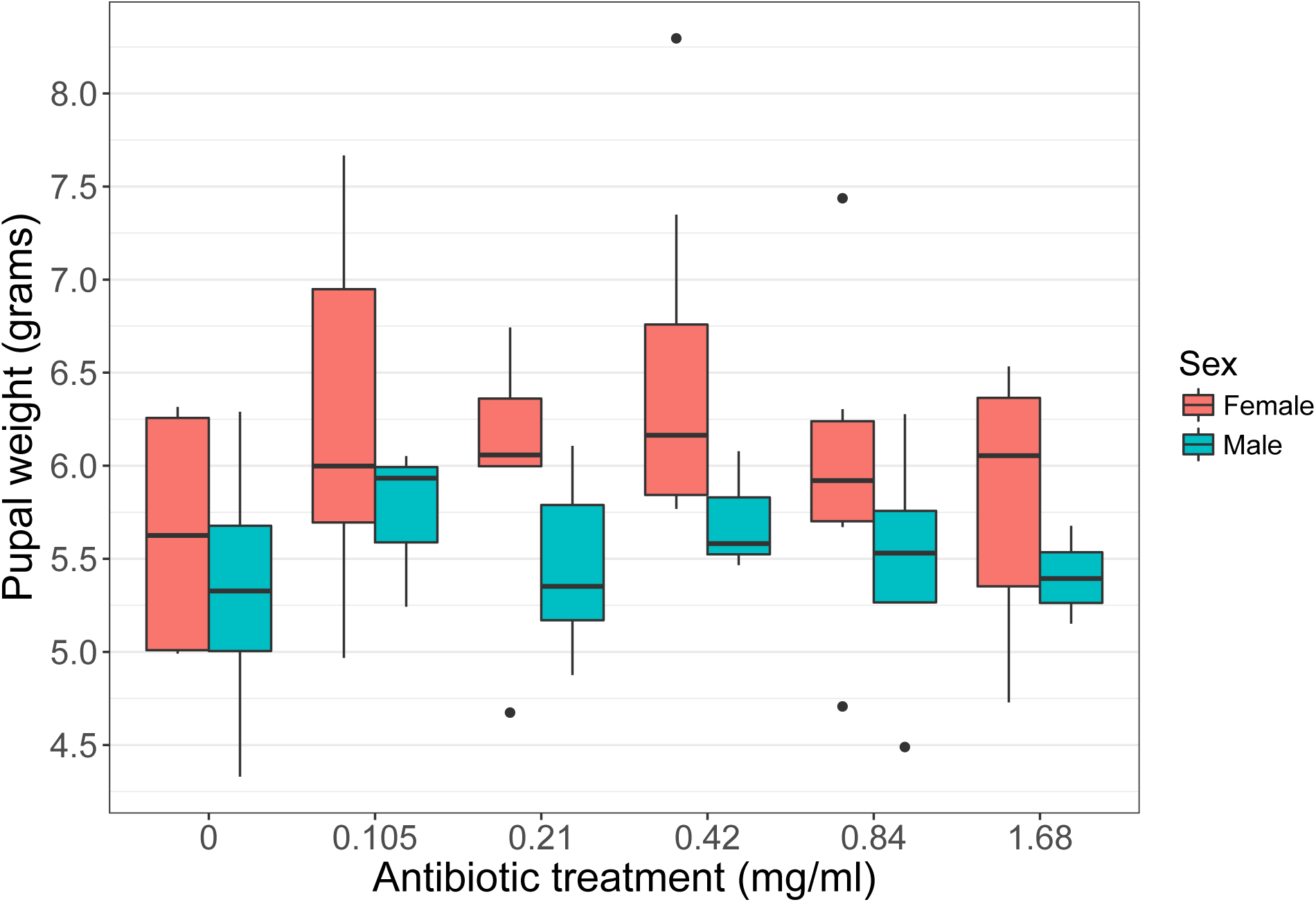
Increasing concentration of an antibiotic cocktail, delivered by a spray applied to *Datura wrightii* leaves prior to feeding, does not reduce *Manduca sexta* growth (N=62). Males and females are plotted separately, as they were expected to differ in size. Fresh weight was measured six days after pupation. Pupal weight correlates with adult fecundity and is often used as a proxy of insect fitness.

## Discussion

Consistent with previous microscopy-based (20–22, 53) and molecular studies (26–29), we found that resident microbial symbionts are generally absent or present only in low numbers in caterpillar guts. As expected for herbivores consuming microbe-rich leaf tissue, diet-derived microbes are transiently present in caterpillar guts, wherein they may be dead or inactive. That the microbial biomass in caterpillar guts is far lower than in the guts or whole bodies of many other animals (Fig. 1A), and also lower than in their food (Fig. 2A), suggests a lack of persistent microbial growth within the gut. Moreover, any potential microbial metabolism might be too limited to substantially affect digestive processes, as illustrated by our observation that *Manduca sexta* caterpillars contain microbial loads orders of magnitude lower than comparably sized animals with resident microbiomes (Fig. S2). In addition to low abundance, the composition of microbes detected in caterpillar guts is highly variable among conspecific individuals (Fig. 1C). Lacking stable populations of core microbial taxa, caterpillar gut microbiomes may be easily influenced by the idiosyncrasies of which microbes are present on a given leaf and in what abundance, and which leaf microbes can survive transit through the digestive tract. Ingested microbes which die within the host may still be beneficial as a food source or by stimulating the immune system, but are not themselves symbionts (following the original definition of symbiosis as the “living together of different species” (referenced in (35)).

Based on the experiment with *M. sexta*, it is unlikely that microbes have cryptic, but essential, functions in caterpillar guts. Antibiotic suppression of viable gut bacterial loads in *M. sexta* had no apparent negative consequences, contrasting sharply with the many examples of major reductions in host growth or survival upon removal of beneficial symbionts (e.g., (54–56)). If anything, caterpillars treated with antibiotics showed slight (but not statistically significant) increases in performance (Fig. 3, Fig. S4B). Antibiotics increase the weight gain of laboratory-bred caterpillars (57–59), and commercially made caterpillar diets often contain antibiotics. This effect, also observed in livestock (60), might reflect microbial parasitism occurring in even apparently healthy caterpillars, and/or costly immune responses to the presence of pathogens (61). Aside from known leaf-specialists, some of the most frequently detected bacterial genera in this study (Table S2), including *Acinetobacter*, *Clostridium*, *Enterobacter*, *Enterococcus*, *Escherichia*, and *Staphylococcus*, have been reported to cause disease in caterpillars under some circumstances (37, 47, 62, 63).

The lack of a resident gut microbiome in caterpillars may directly result from a digestive physiology that is particularly unfavorable to microbial growth (18). The midgut, the largest section of the digestive tract wherein caterpillars digest leaf material and absorb the resulting nutrients (64), is a hostile environment for microbes (24). It is highly alkaline, with pH values often >10 (65) and as high as 12 (66), and contains host-encoded antimicrobial peptides (67). Additional attributes of the caterpillar gut that may hinder microbial colonization include a simple tube-like morphology without obvious microbe-housing structures (18), a continually replaced lining (the peritrophic matrix) covering the midgut epithelium (68) which may prevent biofilm formation, and short retention times (food transit takes ∼2 hours in *M. sexta* (69)). Although some insects harbor symbionts in specialized organs (53), to our knowledge, similar structures have not been reported in caterpillars. Buchner’s foundational survey of animal endosymbiosis describes Lepidoptera only as “a group in which no symbiont bearers have been discovered” ((53), p. 817). Moreover, previous studies did not find abundant microbes outside of the gut (32, 37).

Without the aid of microbial symbionts, how are caterpillars able to overcome the dietary challenges posed by herbivory? First, caterpillars use a combination of mechanical disruption, endogenously produced digestive enzymes, and high pH to extract easily solubilized nutrients, primarily from the contents of plant cells (18, 70, 71). Although this method of processing leaves is relatively inefficient, essential nutrients are not totally absent, so that caterpillars can compensate by simply eating more (18, 64). Some insects likely require microbes for detoxification (16), but many caterpillars possess host-encoded mechanisms for degrading or resisting plant allelochemicals (72). However, there may be a vestigial role for microbes in these processes, as genomes of many Lepidoptera contain microbial genes encoding enzymes with related functions (73, 74). These gene acquisitions may have enabled a symbiont-free feeding strategy.

The caterpillars surveyed here are likely to be representative of most externally leaf-feeding Lepidoptera, as we included a range of families, habitat types, and diet breadths from monophagous to highly generalist. However, a lack of resident gut microbiome in the caterpillar may not apply to the adult butterfly or moth. Compared with larvae, adult butterflies host distinct bacterial communities (32) and high gut microbial loads (75). Many other Lepidoptera lack functioning mouthparts or digestive tracts as adults, and in these groups microbes may be altogether irrelevant to digestion or nutrition. However, we cannot exclude the possibility that microbial symbionts may influence host fitness by their potential activities in eggs or pupae.

The extraordinary diversity and abundance of Lepidoptera (14) indicates that a symbiont-independent feeding strategy can be highly successful. Perhaps such success reflects a release from constraints imposed on other animals that do host and depend on symbionts. There are costs to engaging in mutualisms (e.g., (76–78)), and in a gut microbiome context one cost includes nutrient competition between host and microbes (60). A high availability of food allows caterpillars to “skim the cream” (64), assimilating simple nutrients that might otherwise be used by gut microbes and excreting recalcitrant material. In other words, “Why not do the digestion yourself rather than pay someone else to do it?” ((79), p. 53). Other costs include the risk of gut microbes becoming pathogenic (80, 81), and the potential for pathogens to exploit a gut environment that is hospitable to microbial mutualists. The extreme conditions in the caterpillar midgut may instead exclude all microbial growth, providing some degree of protection against disease.

Dependence on microbes with different physiological tolerances than the host constrains overall niche breadth (7, 77). As compared with groups lacking functional microbiomes, animals whose biology is heavily influenced by microbial mutualists may be less able to switch to new food plants or new habitats over evolutionary time. Indeed, it has been argued that while microbial symbioses can provide novel ecological functions, they may also increase the extinction risk of host lineages (7, 82). As Lepidoptera represent one of the most speciose animal radiations (83), a conspicuous question is whether independence from microbes may, in some cases, facilitate animal diversification.

Caterpillars do not appear to be unique in lacking a resident microbiome that is important for feeding and development. Microbiomes of walking sticks (84), sawfly larvae (85, 86), a saprophagous fly (87), a parasitic horsehair worm (88), a leaf beetle (89, 90), and certain ants (91) display features similar to those we observed in caterpillars. In fact, our data suggest that some vertebrates also have minimal gut microbiomes, and these species may feed relatively autonomously. Feces of the herbivorous brent goose (*Branta bernicla*) had low bacterial loads and a high proportion of plant DNA, and the insectivorous little brown bat (*Myotis lucifugus*) had similarly low fecal bacterial loads (Figs. 1A,B; Table S1). These species exhibit caterpillar-like physiological traits such as a relatively short gut and rapid digestive transit (92, 93). Additional examples in the microbiome literature might be obscured by contaminants masquerading as mutualists (94), a frequent lack of quantitative information (91) and experimental validation of microbial function *in vivo*, and publication bias against “negative results.”

While recent literature has documented extraordinary variation in the types of services provided by microbial symbionts, less explored is variation in the degree to which animals require any such services. Animals likely exist on a spectrum from tightly integrated host-microbe holobionts to simply animals, *sensu stricto*, in which a microbial presence is only relictual (i.e. mitochondria and horizontally transferred genes). Documenting the existence of microbially independent animals as well as their ecological, physiological and phylogenetic contexts, is a first step toward understanding the causes and consequences of evolutionary transitions along this continuum.

## Methods

### Sampling and Sequencing

Caterpillar fecal samples (N=185) were obtained from actively feeding, field-collected individuals in AZ, CO, MA, and NH, USA, and Área de Conservación Guanacaste, Costa Rica. To sample plant microbiomes, we collected leaves from the same branch used to feed caterpillars prior to fecal or gut sampling. Sequence composition was coreelated between feces and guts (*Supplemental Methods*), in line with a previous finding (32). All samples were preserved in 95% ethanol or dry at -20°C (90). We extracted DNA, PCR-amplified the 16S rRNA V4-V5 gene region and sequenced amplicons on an Illumina MiSeq in the same manner as previous insect microbiome studies (32, 90). These DNA extracts and primers were also used for quantitative PCR, which provides microbial biomass estimates concordant with those from microscopy (91) and culturing (Fig. S3B). We did not find evidence that low amplification of caterpillar fecal bacteria is due to primer bias, PCR inhibitors, or storage methods (*Supplemental Methods*).

### Antibiotic Experiment

We collected *Manduca sexta* eggs from *Datura wrightii* plants near Portal, AZ, USA. 72 newly hatched larvae were randomly and evenly divided among six treatments varying from 0–1.68 mg total antibiotics per ml distilled water, and reared in separate unused plastic bags on *D. wrightii* foliage at the Southwestern Research Station. Water with or without antibiotics was sprayed onto leaves, which were briefly dried prior to feeding. The compounds used (rifampicin, tetracycline, streptomycin, in a 1:2:4 ratio) suppressed bacterial symbionts in other insect herbivores (55, 95). We collected a fresh fecal pellet from each caterpillar midway through the final instar, from which one subsample was cultured on LB media, and another used for qPCR and sequencing with the aforementioned protocol. Pupae were weighed six days after pupation and monitored daily for adult eclosion.

### Data Analysis

Statistical analyses were conducted in R (96). Differences in bacterial loads, core sizes, and *M. sexta* performance variables were tested using linear models; residuals were visually inspected for Gaussian structure. The betadisper function in the vegan package was used to compare intraspecific beta diversity. *M. sexta* survival was analyzed using logistic regression. We used a Mantel test to estimate the rank correlation between leaf and fecal microbiome dissimilarities. A Wilcoxon test was used for proportions of plant DNA. Differences in community composition were analyzed using PERMANOVA. DNA sequences, metadata, and R code available at doi:10.6084/m9.figshare.4955648.

## Acknowledgments

We thank the parataxonomists and staff of ACG and The Caterpillar Lab, without whom this project would not have been possible. J Dickerson, K Vaccarello, and H Layton provided invaluable help in the field and laboratory. HA Woods and MD Bowers aided in the planning and interpretation of this study. T Sharpton provided feedback on an earlier version of the manuscript. We also acknowledge administrative support from ACG, the Southwestern Research Station, and A Dietz. TJH was supported by the American Philosophical Society’s Lewis and Clark Fund, the National Science Foundation (NSF) Graduate Research Fellowship Program (1144083) and an NSF Doctoral Dissertation Improvement Grant (1601787). DHJ and WH were supported by the University of Pennsylvania, Wege Foundation, Permian Global, government of Costa Rica, and Guanacaste Dry Forest Conservation Fund.

## Supplemental Methods

**Figure S1.**
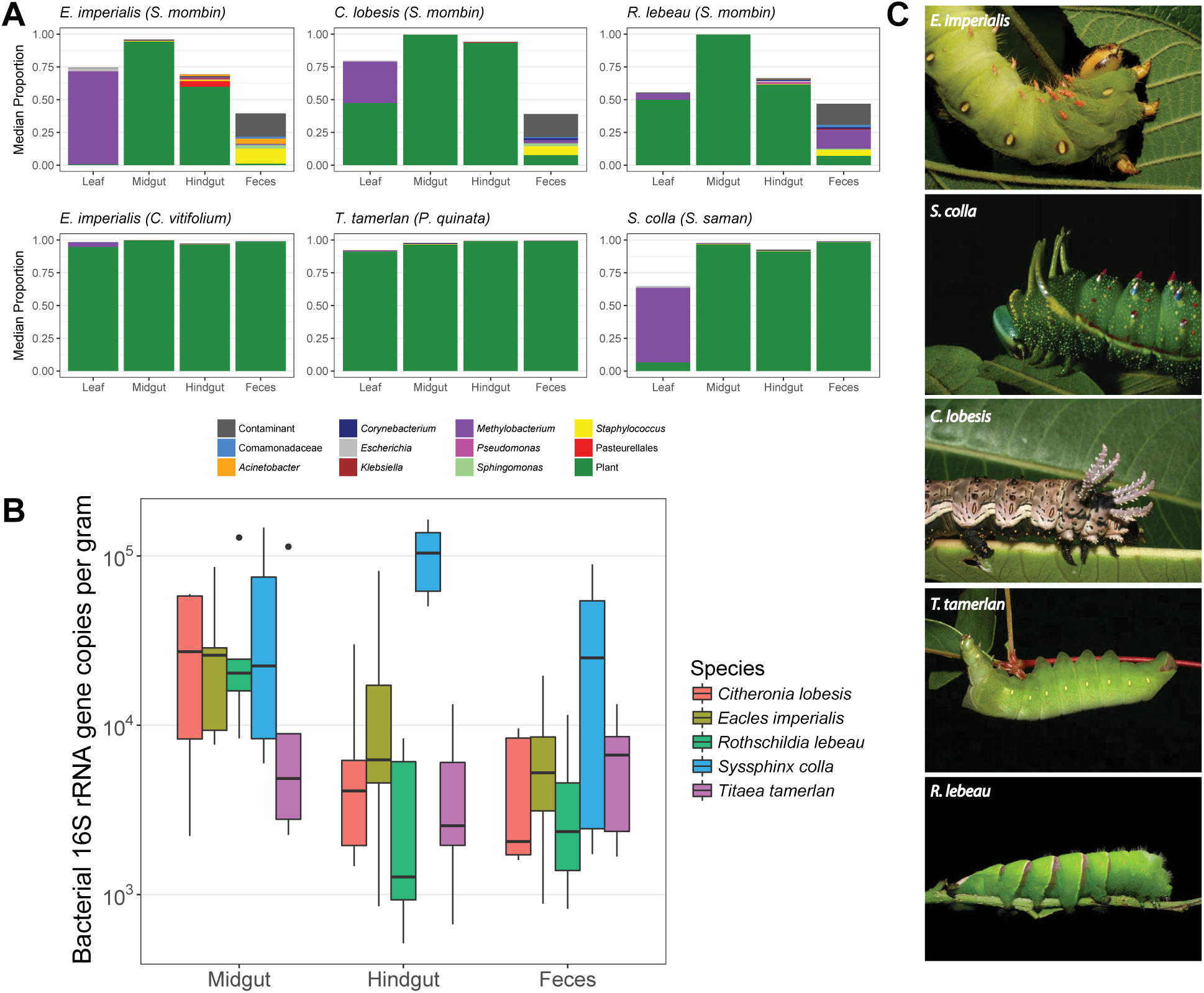
Community composition, bacterial density, and midgut pH in five caterpillar species (Saturniidae) from Área de Conservación Guanacaste (ACG), Costa Rica. A) The composition of sequence libraries from the leaf surface, midgut, hindgut, and feces. The median across five replicate individuals is displayed. The food plant species is indicated in parentheses. Note that one species, *Eacles imperialis*, was reared separately on two plant species. Only plant chloroplast or mitochondrial sequences, reagent contaminants, and the top 10 bacterial genera (among the dissected individuals only) are shown; the remainder of the community (summing to 1) represents sequences from a variety of low-abundance taxa. B) The number of bacterial 16S rRNA gene copies per gram (fresh weight) in homogenized midgut or hindgut tissue and feces (N=5 individuals per species, except *E. imperialis* with 10 individuals (5 each on two plant species)). C) Photographs of each species taken in ACG.

**Figure S2.**
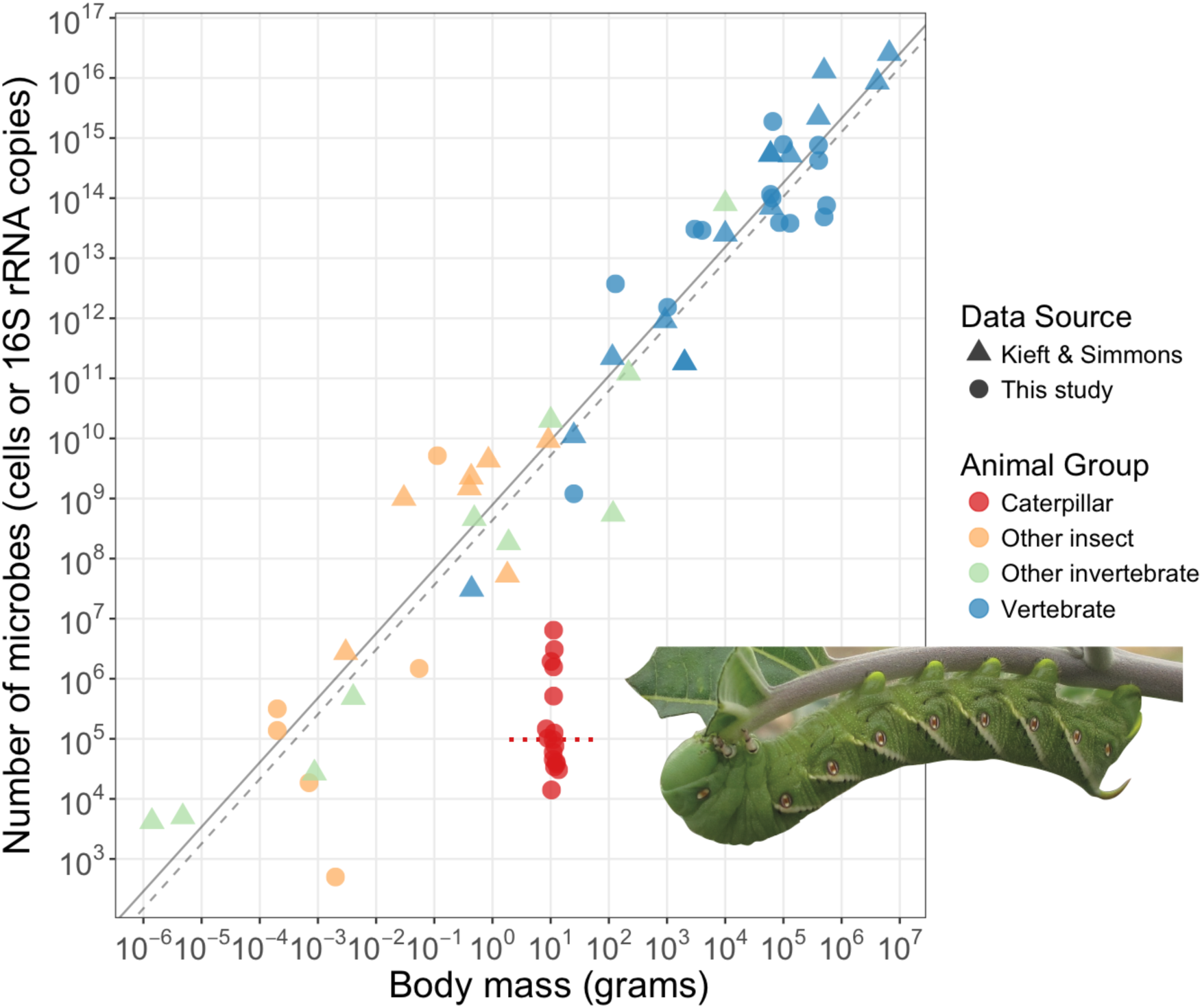
Allometric scaling of whole-individual microbial loads with body size. Triangles and the solid line show data replotted from (50), which were originally measured using microscopy or culturing. Circles show data generated in this study, using quantitative PCR. The dashed regression line is calculated from a model only including non-caterpillar species analyzed in this study, limited to those species with bacterial densities not less than 1/100 of the group median. The red horizontal dotted line indicates the median per-caterpillar bacterial load for 17 *Manduca sexta* individuals collected in Colorado (N=15) or Arizona (N=2). The photograph is *M. sexta* feeding on *D. wrightii*.

**Figure S3.**
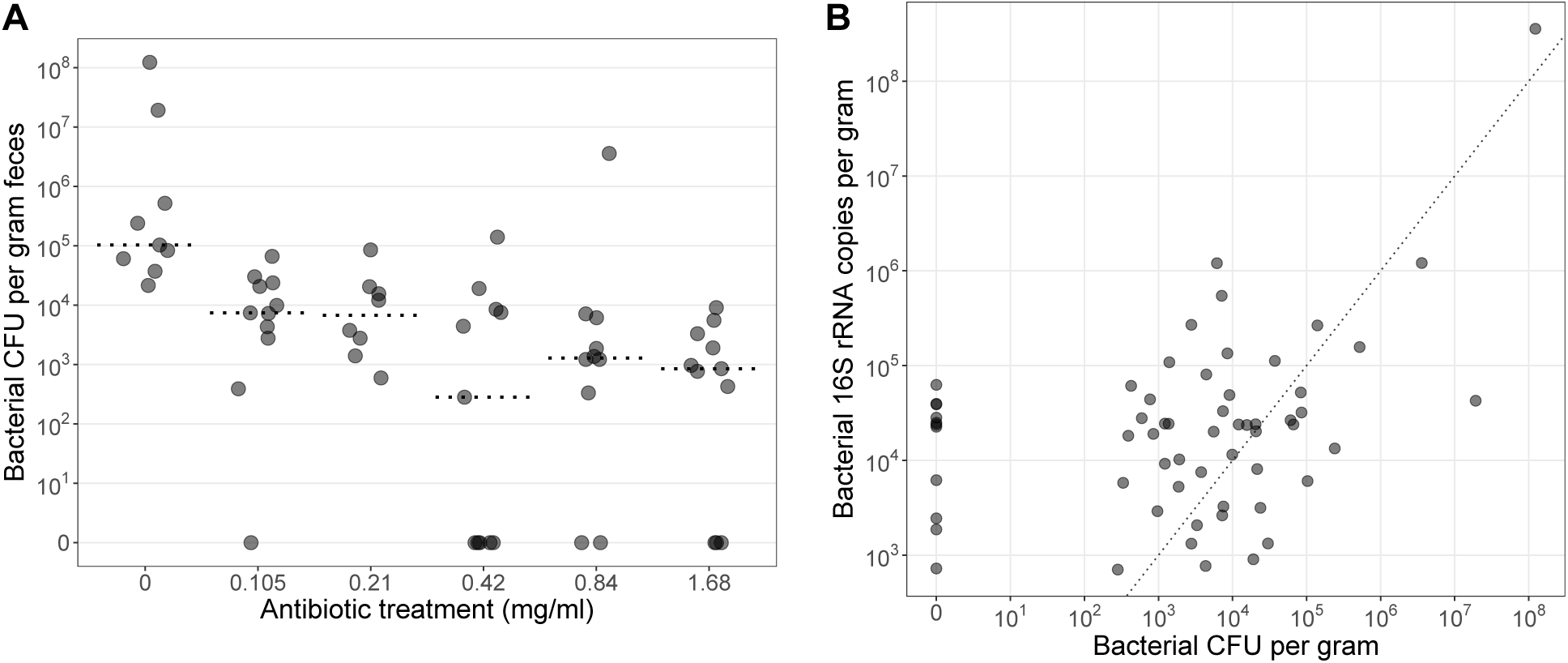
The relationships between antibiotic dose, the number of bacterial colony-forming units cultured on LB media, and the number of bacterial 16S rRNA gene copies measured by qPCR. Fecal samples that yielded no cultured colonies are plotted at 10^0^ on log_10_ axes; for these, nonzero estimates of 16S rRNA gene copies are likely due, in large part, to amplification of DNA from dead or nonviable cells (see *Supplemental Methods*). A) Effect of antibiotic treatment on the number of culturable bacteria in caterpillar feces. Points are individual caterpillars (N=60) and are horizontally jittered for clarity. Dashed lines are medians for each treatment. B) Correlation of bacterial density as measured by culturing versus by DNA quantification. The 1:1 line between the two variables is shown.

**Figure S4.**
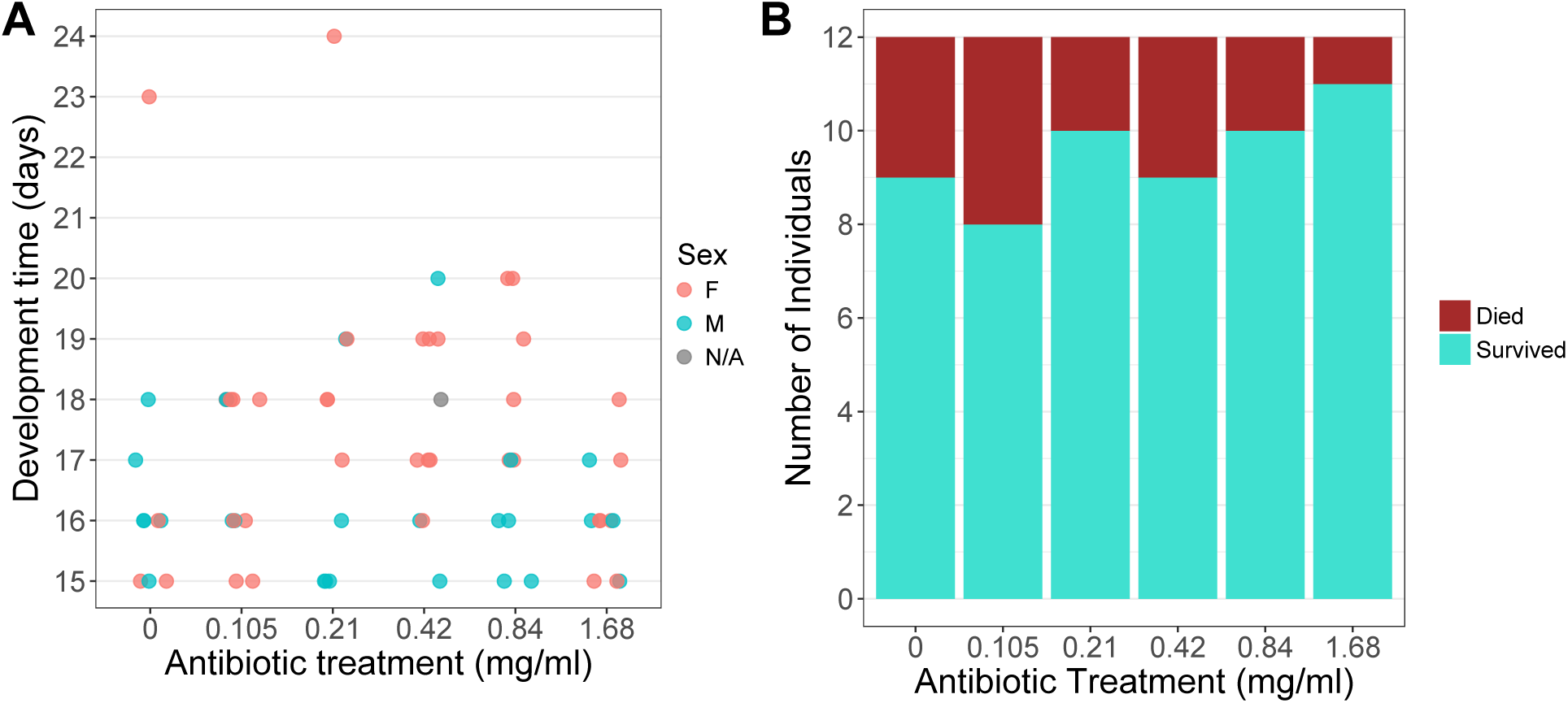
The relationship between antibiotic treatment and other components of *M. sexta* fitness. 12 individuals, randomly selected from the population, were used to initiate each group. A) Number of days from larval hatching from eggs to the cessation of feeding, which marks the beginning of the prepupal stage. Shown are the 64 individuals that survived to this point. We were unable to identify the sex of two individuals. B) The proportion of individuals surviving from larval hatch to adult eclosion, for the control group and each antibiotic treatment.

**Figure S5.**
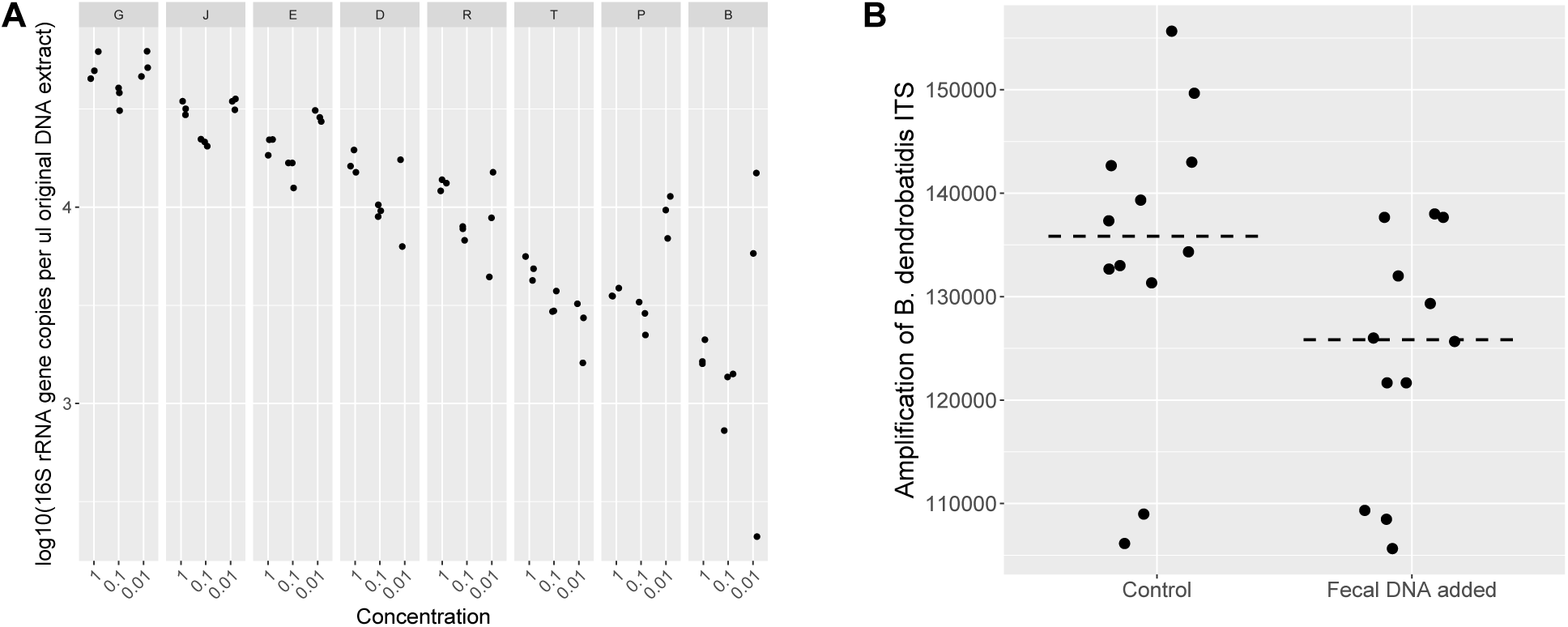
Two tests for PCR inhibitory substances in caterpillar feces. A) Fecal DNA from eight *M. sexta* individuals, arranged left-right by decreasing 16S rRNA gene copy number in original extracts. For each individual, log_10_(16S rRNA gene copies) is shown for the original sample, and for and extracts diluted 1:10 and 1:100 in pure water. Copy number estimates are standardized per ul of original DNA extract. Note that variability between technical replicates increases with low concentrations of template DNA. One sample, D-0.01, had less amplification than negative controls and is not shown. B) Amplification (arbitrary units) of rDNA ITS of *B. dendrobatidis*, a chytrid fungus of amphibians, showing 12 replicate controls (PCR-grade water only) versus 12 reactions to which 5 µl of caterpillar fecal DNA was substituted for water. Means of triplicate reactions are shown. The twelve caterpillar species with the lowest total 16S rRNA gene copy number were used for this test. Dashed lines show medians for each group.

### Sampling

Fecal samples were obtained from wild populations of caterpillars in four regions: Área de Conservación Guanacaste (Costa Rica), New Hampshire and Massachusetts (USA), Boulder County, Colorado (USA), and Portal, Arizona (USA). Caterpillars were collected in ACG under permit #ACG-PI-027-2015 and in Arizona under a Scientific Use Permit from the United States Forest Service. For more details about the ACG landscape and collection, rearing, and identification protocols, see (1–3). Most species were collected as caterpillars, but some ACG specimens were reared from eggs either found on foliage or laid by females caught at light traps (see data file “Metadata_ACG2015”). For some caterpillars we had information on whether they died of parasitoids or disease after sampling, and these samples were discarded in order to focus on apparently healthy individuals. Most caterpillars were sampled in the final or penultimate instar.

All samples were preserved within 30 minutes of defecation, as preliminary evidence suggested rapid (by 6–12 hours) bacterial and fungal growth in excreted fecal pellets, which would render old feces unsuitable as a proxy for gut microbial communities. In five caterpillar species, we did not find evidence for abundant bacterial populations in the midgut (including both ecto- and endoperitrophic spaces) or hindgut that were not captured in feces (Fig. S1A), supporting a previous finding that caterpillar feces approximates the whole-body microbial community (4). Further supporting the use of fresh feces to sample microbes in the caterpillar gut, we found that the inter-individual variation in sequence composition (including nonbacterial DNA) was reflected in fecal samples (Mantel test: midgut *r* = 0.33, *p* = 0.001; hindgut *r* = 0.39, *p* 688 = 0.001).

We preserved gut and fecal samples using either dry storage at -20°C or 95% ethanol (Table S1); both methods are suitable for storing insect microbiome samples and do not substantially alter community composition (5). Approximately 50 mg (fresh weight) of sample was used for DNA extraction. Prior to DNA extraction, ethanol-preserved samples were dried in a vacuum centrifuge; since this also evaporated water, their fresh weight equivalent was estimated using percent water content calculated from *M. sexta* guts or feces. To test whether microbial biomass estimates may have been biased by ethanol storage, we compared PCR amplification for paired ethanol-stored and frozen fecal pellets from eight *M. sexta* individuals. From a collection of pellets defecated by each individual during a 1–2 hour window, separate pellets were randomly chosen for each storage type (note that pre-storage inter-pellet microbial variation is possible even under these relatively controlled conditions). As assessed by a linear mixed-effects model treating individual as a random effect, there was no significant influence of storage method on 16S rRNA gene copy number (χ^2^(1) = 1.09, *p* = 0.30).

For caterpillars in Costa Rica and Colorado, we also sampled microbes from leaves of the same branch as that fed to the caterpillar prior to feces collection. With this strategy we aimed to maximize microbial similarity between the leaves that were sampled and those consumed by the caterpillar, although leaf microbiomes can also vary substantially within a branch (6). These leaves appeared clean and had not, to our knowledge, come into contact with any caterpillars prior to sampling. Leaves from Colorado plants were frozen dry at -20°C and ground under liquid N_2_ with a mortar and pestle prior to DNA extraction (thus including endophytes as well as surface-associated microbes). Leaves from Costa Rican plants were stored in 95% ethanol, and surface-associated microbes were concentrated in a vacuum centrifuge and resuspended in molecular grade water prior to DNA extraction. As this sampling method was not quantitative, we did not perform qPCR on plant samples from Costa Rica and used them only for analyses of microbial composition.

Non-lepidopteran animals were sampled using the same procedures outlined above, with five species preserved in ethanol and 19 preserved dry at -20°C (Table S1). With the exception of two dung beetles feeding on herbivore dung, and the insectivorous bat *M. lucifugus*, these species are either predominantly or exclusively herbivorous, although the type of plant matter consumed (leaves, seeds, fruit, pollen, etc.) varies. We extracted DNA from feces for vertebrates and from subsamples of homogenized whole bodies for insects (as some insects house the majority of symbionts in organs outside the gut). By including all tissue from these insects, we may have underestimated bacterial densities in the particular organs where microbes are housed (Fig. 1A).

### DNA extraction, PCR and sequencing

Following previous studies of insect microbiomes (4, 5, 7), we used the MoBio Powersoil kit to extract DNA (100 µl eluate) from measured amounts of sample material. We then PCR-amplified a portion of the 16S rRNA gene with barcoded 515f/806r primers (8). PCR products were cleaned and normalized (up to 25 ng DNA/sample) using the SequalPrep Normalization kit (Thermo Fisher Scientific), and then sequenced on an Illumina MiSeq. Paired-end sequences of 16S rRNA amplicons were merged, quality-filtered, and clustered into operational taxonomic units (“phylotypes”) at the 97% sequence similarity level using UPARSE (9), and classified using the RDP classifier and Greengenes (10, 11) as previously described (12). The representative sequences of phylotypes unclassified at this stage, and mitochondrial rRNA phylotypes (which could be from plant, insect, fungal or other mitochondria) were aligned to the NCBI nonredundant nucleotide database (nt) using BLAST for taxonomic identification. (Many universal 16S primers amplify rRNA genes of chloroplasts and mitochondria as well as bacteria and archaea (13)).

As bacterial DNA is ubiquitous in laboratory reagents used for DNA extraction and PCR, and especially problematic with low-biomass samples (such as caterpillar feces) (14), we removed contaminants from our samples using information from the 22 DNA extraction blanks and PCR no-template controls that yielded >100 bacterial sequences. Importantly, phylotypes detected in these blanks are not exclusively composed of reagent contaminants, because they receive some input from sample DNA during laboratory processing (15). As high-biomass samples are both least likely to experience reagent contamination (14), and themselves most likely to be the source of “real” sample phylotypes identified in blanks, they can be used to distinguish between laboratory contaminants and true sample sequences (15). We classified contaminants as phylotypes present at ≥1% abundance in one or more blank samples, excepting phylotypes present at ≥1% abundance in one or more of the best-amplifying samples (the top third in 16S rRNA gene copy number as measured by qPCR). These 25 phylotypes were removed from the dataset prior to analyses of bacterial abundance and composition (they are retained only in Fig. S1A). This approach does not include other types of contaminants introduced prior to DNA extraction, such as those from human skin. Finally, we note that the high relatedness between microbes commonly present in laboratory reagents (listed in (14)) and those present in soil, water and leaves—all possible genuine microbial inputs to the caterpillar gut—precludes a taxonomy-based approach to removing contaminants.

### Sequence Data Analysis

All analyses were conducted in R version 3.3.2 and are available in the file “Hammer2017_Rcode.R”. Analyses involving bacterial composition were limited to samples with at least 100 bacterial sequences. To calculate phylotype-level overlap between fecal and plant samples, “phylotypes detected on leaves” are defined as those present at any abundance in any plant sample in our dataset. New England and Arizona fecal samples which lack paired plant samples were excluded from this comparison. In measuring intraspecific beta diversity and core microbiome size in caterpillars and other animals, we excluded species with fewer than three replicate individuals. Further, to be conservative, only caterpillars sampled from the same location, and feeding on the same species of plant were compared. As the number of replicates could affect these metrics, and varied among species, we iterated these analyses over multiple combinations of only three replicates per species.

### Quantitative PCR

We measured 16S rRNA gene copy number using quantitative PCR with the same primers and DNA extracts as above. Reaction conditions and other details are specified in (16). Each sample was run in triplicate (except 11 non-caterpillar species for which limited DNA was available, which were run singly) and the mean of these technical replicates was used for subsequent analyses. Standard curves were calculated using purified genomic DNA from *E. coli* DH10B, which has seven 16S rRNA operons per genome (17). The median copy number of 31 qPCR’d DNA extraction blanks was subtracted from sample copy numbers. Resulting counts of total 16S rRNA genes in samples were then multiplied by the proportion of noncontaminant bacterial sequences identified from the same DNA extract, resulting in estimates of bacterial 16S copy numbers.

It is unlikely that the low amplification we found in caterpillar samples results from primer bias against abundant bacterial taxa. First, these primers successfully amplified bacteria in non-lepidopteran animals, even when in some cases (such as aphids, (18)), the dominant symbiont has been strictly vertically transmitted between hosts for tens of millions of years. Even in this case, divergence from free-living relatives has not been so great that its 16S rRNA gene is un-amplifiable using 515f/806r primers. Second, the caterpillar gut-associated microbial taxa we found are similar to those reported as being relatively (i.e., in terms of the proportion of sequence libraries) abundant in metagenomic surveys (19, 20) and amplicon-based studies using different 16S rRNA-targeting primer pairs (e.g., (21–25)).

To estimate the relationship between body size and whole-animal microbial loads (Fig. S2), we combined published data from (26) with body mass data we calculated directly or derived from other studies (see supplemental file “Body_mass_data.txt”). To restrict the allometric scaling relationship for noncaterpillar animals to those species likely to harbor resident microbiomes, we removed species that had bacterial densities < 1/100^th^ of the group median. These species were the goose *Branta bernicla*, the bat *Myotis lucifugus*, and the dung beetle *Geotrupes stercorosus*. The body size of two *M. sexta* individuals from Arizona was not recorded and so we substituted the median from other *M. sexta*. Furthermore, as we only had direct gut mass measurements for *M. sexta* (30–40% of body mass), for species sampled using feces (including *M. sexta*) we calculated total microbial loads by multiplying 16S rRNA gene density in feces by body mass. This procedure is likely to have slightly overestimated the microbial load for these species. Despite the numerous methodological uncertainties, microbial counts from (26) and our qPCR-based data, and their allometric scaling relationship with body size (excepting *M. sexta*) were remarkably similar (compare solid and dashed line in Fig. S2).

### PCR inhibition assays

To examine whether low 16S rRNA gene copy number estimates in caterpillar samples are an artifact of caterpillar-specific PCR inhibitors, we used two distinct approaches. First, we tested whether diluting extracted DNA improves PCR amplification by minimizing inhibitor effects (27). However, 1:10 and 1:100 dilutions of fecal DNA from eight *M. sexta* individuals did not have this effect (Fig. S5A). Second, we individually added the twelve lowest-amplifying caterpillar fecal samples—which might be especially likely to contain PCR inhibitors—to qPCR reactions with targeted primers and a template highly unlikely to be present in caterpillar feces (rDNA ITS region of *Batrachochytrium dendrobatidis* strain JEL270, a chytrid fungus pathogenic to amphibians). As compared to replicate reactions with pure molecular-grade water, adding caterpillar fecal DNA reduced amplification of *B. dendrobatidis* rDNA by 7.4% (Fig. S5B). This inhibition effect, which is also present in feces of humans (27) and likely many other species, is miniscule relative to the difference in bacterial loads between caterpillars and non-lepidopterans spanning multiple orders of magnitude (Fig. 1A). Therefore, the relatively low PCR amplification of 16S rRNA genes from caterpillar feces is most likely due to low microbial biomass rather than high PCR-inhibitory substances.

### Additional information on the antibiotic experiment

*M. sexta* larval feces production was measured by collecting, drying (50°C for 24 hours), and weighing all fecal pellets in the final instar. To culture bacteria, we plated a dilution series (in sterilized phosphate-buffered saline) of weighed (10–20 mg) subsamples of feces, incubated in aerobic conditions at 37°C. After 24 hours, visible colonies were counted and then, if present, collected *en masse* from the agar surface for sequencing using a sterile swab. This plate-scrape method produces a list of the most abundant bacterial phylotypes potentially culturable using our approach. It should be noted that the presence of fecal bacteria in culture demonstrates that these taxa were viable, but not necessarily growing or metabolically active, while in the caterpillar gut.

### Comparison of biomass estimates and evidence of extracellular DNA

Among *M. sexta* fecal samples collected during the antibiotic experiment, we found that qPCR-estimated bacterial abundances were correlated with the number of cultured bacterial colonies (see Results; Fig. S3B). Eleven individuals’ fecal pellets did not produce any bacterial colonies whatsoever, but did contain measurable levels of DNA (Fig. S3B), and excluding these “zero-colony” samples yielded a stronger association between bacterial colony counts and 16S rRNA gene copy number (*r* = 0.51, *p* = 0.0002). This result could stem from the presence of bacteria that cannot grow aerobically or on LB. Alternatively, it may be due to PCR amplification of extracellular DNA or DNA from dead or otherwise nonviable cells (16). To evaluate these possibilities, we compared the phylotypes (identified by 16S rRNA gene sequencing) in zero-colony fecal samples to those from other samples that did yield colonies, in which bacterial biomass was swabbed directly from the agar surface and sequenced. Most of the 16S rRNA gene sequences in the zero-colony fecal samples (median 84%, interquartile range: 74–95%) belong to phylotypes cultured from other samples, suggesting that qPCR may have overestimated viable bacterial loads by amplifying DNA from lysed or nonviable cells. If the fraction of the gut microbiome originating from dead or nonviable cells is disproportionately high in caterpillars in general (e.g., due to their digestive physiology – see Discussion), then the difference in living, active microbial biomass between caterpillars and other animals (Fig. 1A) may have been underestimated.

## References

1. McFall-Ngai M, et al. (2013) Animals in a bacterial world, a new imperative for the life sciences. Proc Natl Acad Sci 110(9):3229–3236.

2. Sudakaran S, Kost C, Kaltenpoth M (2017) Symbiont acquisition and replacement as a source of ecological innovation. Trends Microbiol 25(5):375–390.

3. Janson EM, Stireman JO, Singer MS, Abbot P (2008) Phytophagous insect-microbe mutualisms and adaptive evolutionary diversification. Evolution 62(5):997–1012.

4. Frago E, Dicke M, Godfray HCJ (2012) Insect symbionts as hidden players in insect–plant interactions. Trends Ecol Evol 27(12):705–711.

5. Sommer F, Bäckhed F (2013) The gut microbiota — masters of host development and physiology. Nat Rev Microbiol 11(4):227–38.

6. Douglas AE (2014) Symbiosis as a general principle in eukaryotic evolution. Cold Spring Harb Perspect Biol 6(2):1–14.

7. Moran NA (2002) The ubiquitous and varied role of infection in the lives of animals and plants. Am Nat 160:S1–8.

8. Zilber-Rosenberg I, Rosenberg E (2008) Role of microorganisms in the evolution of animals and plants: The hologenome theory of evolution. FEMS Microbiol Rev 32(5):723–735.

9. Bordenstein SR, Theis KR (2015) Host biology in light of the microbiome: Ten principles of holobionts and hologenomes. PLoS Biol 13(8):e1002226.

10. Gilbert SF, Sapp J, Tauber AI (2012) A symbiotic view of life: We have never been individuals. Q Rev Biol 87(4):325–341.

11. Lederberg J, McCray AT (2001) ‘Ome Sweet ’Omics—a genealogical treasury of words. Sci:8.

12. Russell JA, Dubilier N, Rudgers JA (2014) Nature’s microbiome: Introduction. Mol Ecol 23(6):1225–1237.

13. Vavre F, Kremer N (2014) Microbial impacts on insect evolutionary diversification: from patterns to mechanisms. Curr Opin Insect Sci 4(1):29–34.

14. Scoble MJ (1992) The Lepidoptera (Oxford University Press).

15. Stamp NE, Casey TM (1993) Caterpillars: ecological and evolutionary constraints on foraging (Chapman & Hall Ltd.).

16. Hammer TJ, Bowers MD (2015) Gut microbes may facilitate insect herbivory of chemically defended plants. Oecologia 179(1):1–14.

17. Douglas AE (2009) The microbial dimension in insect nutritional ecology. Funct Ecol 23(1):38–47.

18. Appel H (1994) The chewing herbivore gut lumen: physicochemical conditions and their impact on plant nutrients, allelochemicals, and insect pathogens. Insect–Plant Interactions, ed Bernays EA, pp 209–223.

19. Bernays EA, Janzen DH (1988) Saturniid and sphingid caterpillars: Two ways to eat leaves. Ecology 69(4):1153–1160.

20. Shannon AL, Attwood G, Hopcroft DH, Christeller JT (2001) Characterization of lactic acid bacteria in the larval midgut of the keratinophagous lepidopteran, *Hofmannophila pseudospretella*. Lett Appl Microbiol 32(1):36–41.

21. Kukal O, Dawson TE (1989) Temperature and food quality influences feeding behavior, assimilation efficiency and growth rate of arctic woolly-bear caterpillars. Oecologia 79(4):526–532.

22. Vilanova C, Latorre A, Porcar M (2016) The generalist inside the specialist: Gut bacterial communities of two insect species feeding on toxic plants are dominated by *Enterococcus* sp. Front Microbiol 7(June):1–8.

23. Anand AAP, et al. (2010) Isolation and characterization of bacteria from the gut of *Bombyx mori* that degrade cellulose, xylan, pectin and starch and their impact on digestion. J Insect Sci 10(107):1–20.

24. Broderick NA, Raffa KF, Goodman RM, Handelsman J (2004) Census of the bacterial community of the gypsy moth larval midgut by using culturing and culture-independent methods. Appl Environ Microbiol 70(1):293–300.

25. Kingsley VV. (1972) Persistence of intestinal bacteria in the developmental stages of the monarch butterfly (*Danaus plexippus*). J Invertebr Pathol 20:51–58.

26. Priya NG, Ojha A, Kajla MK, Raj A, Rajagopal R (2012) Host plant induced variation in gut bacteria of *Helicoverpa armigera*. PLoS One 7(1):e30768.

27. Staudacher H, et al. (2016) Variability of bacterial communities in the moth *Heliothis virescens* indicates transient association with the host. PLoS One 11(5):1–21.

28. Mason CJ, Raffa KF (2014) Acquisition and structuring of midgut bacterial communities in gypsy moth (Lepidoptera: Erebidae) larvae. Environ Entomol 43(3):595–604.

29. Whitaker M, Pierce N, Salzman S, Kaltenpoth M, Sanders J (2016) Microbial communities of lycaenid butterflies do not correlate with larval diet. Front Microbiol 7:1920.

30. Salter SJ, et al. (2014) Reagent and laboratory contamination can critically impact sequence-based microbiome analyses. BMC Biol 12(1):87.

31. Carini P, et al. (2016) Relic DNA is abundant in soil and obscures estimates of soil microbial diversity. Nat Microbiol 53(9):680840.

32. Hammer TJ, McMillan WO, Fierer N (2014) Metamorphosis of a butterfly-associated bacterial community. PLoS One 9(1):e86995.

33. Lighthart B (1988) Some changes in gut bacterial flora of field-grown *Peridroma saucia* (Lepidoptera: Noctuidae) when brought into the laboratory. Appl Environ Microbiol 54(7):1896–8.

34. Berg RD (1996) The indigenous gastrointestinal microflora. Trends Microbiol 4(11):430–435.

35. Douglas AE (1994) Symbiotic Interactions (Oxford University Press).

36. Steinhaus EA (1956) Microbial control—the emergence of an idea doi:10.1017/CBO9781107415324.004.

37. Bucher GE (1967) Pathogens of tobacco and tomato hornworms. J Invertebr Pathol 9:82–89.

38. Ehrlich P, Raven P (1964) Butterflies and plants: a study in coevolution. Evolution 18(4):586–608.

39. I-M-Arnold A, et al. (2016) Forest defoliator pests alter carbon and nitrogen cycles. R Soc Open Sci 3(10):160361.

40. Pearse IS, Altermatt F (2013) Predicting novel trophic interactions in a non-native world. Ecol Lett 16(8):1088–94.

41. Thomas CD, et al. (1987) Incorporation of a European weed into the diet of a North American herbivore. Evolution 41(4):892–901.

42. Gibson CM, Hunter MS (2010) Extraordinarily widespread and fantastically complex: comparative biology of endosymbiotic bacterial and fungal mutualists of insects. Ecol Lett 13(2):223–34.

43. Falony G, et al. (2016) Population-level analysis of gut microbiome variation. Science 352(6285):560–564.

44. Moran NA, Hansen AK, Powell JE, Sabree ZL (2012) Distinctive gut microbiota of honey bees assessed using deep sampling from individual worker bees. PLoS One 7(4):e36393.

45. Sudakaran S, Salem H, Kost C, Kaltenpoth M (2012) Geographical and ecological stability of the symbiotic mid-gut microbiota in European firebugs, *Pyrrhocoris apterus* (Hemiptera, Pyrrhocoridae). Mol Ecol 21:6134–6151.

46. Vorholt JA (2012) Microbial life in the phyllosphere. Nat Rev Microbiol 10(12):828–840.

47. Haloi K, Kankana M, Nath R, Devi D (2016) Characterization and pathogenicity assessment of gut-associated microbes of muga silkworm *Antheraea assamensis* Helfer (Lepidoptera: Saturniidae). J Invertebr Pathol 138:73–85.

48. Grice EA, et al. (2009) Topographical and temporal diversity of the human skin microbiome. Science 324(5931):1190–2.

49. Brooks AW, Kohl KD, Brucker RM, van Opstal EJ, Bordenstein SR (2016) Phylosymbiosis: Relationships and functional effects of microbial communities across host evolutionary history. PLoS Biol 14(11):e2000225.

50. Kieft TL, Simmons KA (2015) Allometry of animal-microbe interactions and global census of animal-associated microbes. Proc R Soc B 282:20150702.

51. Honek A (1993) Intraspecific variation in body size and fecundity in insects: a general relationship. Oikos 66(3):483–492.

52. Stillwell RC, Davidowitz G (2010) A developmental perspective on the evolution of sexual size dimorphism of a moth. Proc R Soc B 277(1690):2069–2074.

53. Buchner P (1965) Endosymbiosis of animals with plant microorganisms (John Wiley & Sons).

54. Salem H, Kreutzer E, Sudakaran S, Kaltenpoth M (2012) Actinobacteria as essential symbionts in firebugs and cotton stainers (Hemiptera, Pyrrhocoridae). Environ Microbiol 15:1956–1968.

55. Ceja-Navarro JA, et al. (2015) Gut microbiota mediate caffeine detoxification in the primary insect pest of coffee. Nat Commun 6:7618.

56. Raymann K, Shaffer Z, Moran NA (2017) Antibiotic exposure perturbs the gut microbiota and elevates mortality in honeybees. PLoS Biol 15(3):e2001861.

57. van der Hoeven R, Betrabet G, Forst S (2008) Characterization of the gut bacterial community in Manduca sexta and effect of antibiotics on bacterial diversity and nematode reproduction. FEMS Microbiol Lett 286(2):249–56.

58. Murthy MR, Sreenivasaya M (1953) Effect of antibiotics on the growth of the silkworm *Bombyx mori* L. Nature 172(4380):684–685.

59. Visôtto LE, Oliveira MGA, Guedes RNC, Ribon AOB, Good-God PIV (2009) Contribution of gut bacteria to digestion and development of the velvetbean caterpillar, *Anticarsia gemmatalis*. J Insect Physiol 55(3):185–91.

60. Gaskins HR, Collier CT, Anderson DB (2002) Antibiotics as growth promotants: mode of action. Anim Biotechnol 13:29–42.

61. Rolff J, Siva-Jothy MT, Rolff M. T. J. S-J (2003) Invertebrate ecological immunology. Science 301(5632):472–475.

62. Bulla LA, Rhodes RA, St. Julian G (1975) Bacteria as insect pathogens. Annu Rev Microbiol 29:163–190.

63. Peleg AY, et al. (2009) Galleria mellonella as a model system to study Acinetobacter baumannii pathogenesis and therapeutics. Antimicrob Agents Chemother 53(6):2605–2609.

64. Dow JAT (1986) Insect midgut function. Adv In Insect Phys 19:187–328.

65. Johnson K, Felton G (1996) Potential influence of midgut pH and redox potential on protein utilization in insect herbivores. Arch Insect Biochem Physiol 32:85–105.

66. Dow JAT (1984) Extremely high pH in biological systems: a model for carbonate transport. Am J Physiol 246:R633–R636.

67. Jiang H, Vilcinskas A, Kanost MR (2010) Immunity in Lepidopteran insects. Invertebrate Immunity, ed Söderhäll K (Springer), pp 181–204.

68. Hegedus D, Erlandson M, Gillott C, Toprak U (2009) New insights into peritrophic matrix synthesis, architecture, and function. Annu Rev Entomol 54:285–302.

69. Brinkmann N, Tebbe CC (2007) Leaf-feeding larvae of *Manduca sexta* (Insecta, Lepidoptera) drastically reduce copy numbers of aadA antibiotic resistance genes from transplastomic tobacco but maintain intact aadA genes in their feces. Environ Biosafety Res 6:121–133.

70. Santos CD, Ferreira C, Terra WR (1983) Consumption of food and spatial organization of digestion in the cassava hornworm, *Erinnyis ello*. J Insect Physiol 29(9):707–714.

71. Barbehenn R V. (1992) Digestion of uncrushed leaf tissues by leaf-snipping larval Lepidoptera. Oecologia 89(2):229–235.

72. Després L, David J-P, Gallet C (2007) The evolutionary ecology of insect resistance to plant chemicals. Trends Ecol Evol 22(6):298–307.

73. Sun BF, et al. (2013) Multiple ancient horizontal gene transfers and duplications in lepidopteran species. Insect Mol Biol 22(1):72–87.

74. Wybouw N, et al. (2014) A gene horizontally transferred from bacteria protects arthropods from host plant cyanide poisoning. eLife 3:e02365.

75. Ravenscraft A, Berry M, Hammer T, Peay K, Boggs C (2017) Structure and function of the bacterial and fungal gut flora of Neotropical butterflies. bioRxiv. doi:10.1101/128884.

76. Frederickson ME, et al. (2012) The direct and ecological costs of an ant-plant symbiosis. Am Nat 179(6):768–78.

77. Nougué O, Gallet R, Chevin L-M, Lenormand T (2015) Niche limits of symbiotic gut microbiota constrain the salinity tolerance of brine shrimp. Am Nat 186(3):390–403.

78. Simonsen AK, Dinnage R, Barrett LG, Prober SM, Thrall PH (2017) Symbiosis limits establishment of legumes outside their native range at a global scale. Nat Commun 8:14790.

79. Janzen DH (1985) The natural history of mutualisms. The Biology of Mutualism, ed Boucher DH (Oxford University Press), pp 40–99.

80. Broderick NA, et al. (2009) Contributions of gut bacteria to *Bacillus thuringiensis*-induced mortality vary across a range of Lepidoptera. BMC Biol 7:11.

81. Young BC, et al. (2017) Severe infections emerge from the microbiome by adaptive evolution. bioRxiv. doi:10.1101/116681.

82. Bennett GM, Moran NA (2015) Heritable symbiosis: The advantages and perils of an evolutionary rabbit hole. Proc Natl Acad Sci 112(33):10169–10176.

83. Wiens JJ, Lapoint RT, Whiteman NK (2015) Herbivory increases diversification across insect clades. Nat Commun 6:1–7.

84. Shelomi M, Lo W-S, Kimsey LS, Kuo C-H (2013) Analysis of the gut microbiota of walking sticks (Phasmatodea). BMC Res Notes 6(1):368.

85. Lucarotti CJ, Whittome-Waygood BH, Levin DB (2011) Histology of the larval *Neodiprion abietis* (Hymenoptera: Diprionidae) digestive tract. Psyche:1–10.

86. Whittome B, Graham RI, Levin DB (2007) Preliminary examination of gut bacteria from *Neodiprion abietis* (Hymenoptera: Diprionidae) larvae. J Entomol Soc Ontario 138:49–63.

87. Šustr V, Stingl U, Brune A (2014) Microprofiles of oxygen, redox potential, and pH, and microbial fermentation products in the highly alkaline gut of the saprophagous larva of *Penthetria holosericea* (Diptera: Bibionidae). J Insect Physiol 67:64–69.

88. Hudson AJ, Floate KD (2009) Further evidence for the absence of bacteria in horsehair worms (Nematomorpha: Gordiidae). J Parasitol 95(6):1545–1547.

89. Taylor EC (1985) Cellulose digestion in a leaf eating insect, the mexican bean beetle, *Epilachna varivestis*. Insect Biochem 15(2):315–320.

90. Hammer TJ, Dickerson JC, Fierer N (2015) Evidence-based recommendations on storing and handling specimens for analyses of insect microbiota. PeerJ 3:e1190.

91. Sanders J, et al. (2017) Dramatic differences in gut bacterial densities help to explain the relationship between diet and habitat in rainforest ants. bioRxiv. doi:10.1101/114512.

92. Buchler ER (1975) Food transit time in *Myotis lucifugus* (Chiroptera: Vespertilionidae). J Mammal 56(1):252–255.

93. Buchsbaum R, Wilson J, Valiela I (1986) Digestibility of plant constitutents by Canada Geese and Atlantic Brant. Ecology 67(2):386–393.

94. Lauder AP, et al. (2016) Comparison of placenta samples with contamination controls does not provide evidence for a distinct placenta microbiota. Microbiome 4(1):29.

95. Wilkinson TL (1998) The elimination of intracellular microorganisms from insects: An analysis of antibiotic-treatment in the pea aphid (*Acyrthosiphon pisum*). Comp Biochem Physiol Part A 119:871–881.

96. R Core Team (2016) R: A language and environment for statistical computing. R Found Stat Comput. Available at: http://www.r-project.org.

## References for Supplemental Methods

1. Janzen DH, Hallwachs W (2011) Joining inventory by parataxonomists with DNA barcoding of a large complex tropical conserved wildland in northwestern Costa Rica. PLoS One 6(8):e18123.

2. Janzen DH, et al. (2009) Integration of DNA barcoding into an ongoing inventory of complex tropical biodiversity. Mol Ecol Resour 9:1–26.

3. Janzen DH, Hallwachs W (2016) DNA barcoding the Lepidoptera inventory of a large complex tropical conserved wildland, Area de Conservacion Guanacaste, northwestern Costa Rica. Genome 59(9):641–660.

4. Hammer TJ, McMillan WO, Fierer N (2014) Metamorphosis of a butterfly-associated bacterial community. PLoS One 9(1):e86995.

5. Hammer TJ, Dickerson JC, Fierer N (2015) Evidence-based recommendations on storing and handling specimens for analyses of insect microbiota. PeerJ 3:e1190.

6. Espinosa-Garcia FJ, Langenheim JH (1990) The endophytic fungal community in leaves of a coastal redwood population diversity and spatial patterns. New Phytol 116(1):89–97.

7. Hammer TJ, et al. (2016) Treating cattle with antibiotics affects greenhouse gas emissions, and microbiota in dung and dung beetles. Proc R Soc B 283(1831):20160150.

8. Caporaso JG, et al. (2012) Ultra-high-throughput microbial community analysis on the Illumina HiSeq and MiSeq platforms. ISME J 6(8):1621–1624.

9. Edgar RC (2013) UPARSE: highly accurate OTU sequences from microbial amplicon reads. Nat Methods 10(10):996–8.

10. Wang Q, Garrity GM, Tiedje JM, Cole JR (2007) Naive Bayesian classifier for rapid assignment of rRNA sequences into the new bacterial taxonomy. Appl Environ Microbiol 73(16):5261–5267.

11. McDonald D, et al. (2012) An improved Greengenes taxonomy with explicit ranks for ecological and evolutionary analyses of bacteria and archaea. ISME J 6(3):610–618.

12. Ramirez KS, et al. (2014) Biogeographic patterns in below-ground diversity in New York City’s Central Park are similar to those observed globally. Proc R Soc B Biol Sci 281:20141988.

13. Rastogi G, Tech JJ, Coaker GL, Leveau JHJ (2010) A PCR-based toolbox for the culture-independent quantification of total bacterial abundances in plant environments. J Microbiol Methods 83(2):127–132.

14. Salter SJ, et al. (2014) Reagent and laboratory contamination can critically impact sequence-based microbiome analyses. BMC Biol 12(1):87.

15. Lazarevic V, Gaïa N, Girard M, Schrenzel J (2016) Decontamination of 16S rRNA gene amplicon sequence datasets based on bacterial load assessment by qPCR. BMC Microbiol 16(1):73.

16. Carini P, et al. (2016) Relic DNA is abundant in soil and obscures estimates of soil microbial diversity. Nat Microbiol 53(9):680840.

17. Durfee T, et al. (2008) The complete genome sequence of *Escherichia coli* DH10B: Insights into the biology of a laboratory workhorse. J Bacteriol 190(7):2597–2606.

18. Clark MA, Moran NA, Baumann P (1999) Sequence evolution in bacterial endosymbionts having extreme base compositions. Mol Biol Evol 16:1586–1598.

19. Belda E, et al. (2011) Microbial diversity in the midguts of field and lab-reared populations of the European corn borer *Ostrinia nubilalis*. PLoS One 6(6):e21751.

20. Xia X, et al. (2017) Metagenomic sequencing of diamondback moth gut microbiome unveils key holobiont adaptations for herbivory. Front Microbiol. doi:10.3389/fmicb.2017.00663.

21. Pinto-Tomás AA, et al. (2011) Comparison of midgut bacterial diversity in tropical caterpillars (Lepidoptera: Saturniidae) fed on different diets. Environ Entomol 40(5):1111–1122.

22. Broderick NA, Raffa KF, Goodman RM, Handelsman J (2004) Census of the bacterial community of the gypsy moth larval midgut by using culturing and culture-independent methods. Appl Environ Microbiol 70(1):293–300.

23. Brinkmann N, Martens R, Tebbe CC (2008) Origin and diversity of metabolically active gut bacteria from laboratory-bred larvae of *Manduca sexta* (Sphingidae, Lepidoptera, Insecta). Appl Environ Microbiol 74(23):7189–7196.

24. Staudacher H, et al. (2016) Variability of bacterial communities in the moth *Heliothis virescens* indicates transient association with the host. PLoS One 11(5):1–21.

25. Mason CJ, Raffa KF (2014) Acquisition and structuring of midgut bacterial communities in gypsy moth (Lepidoptera: Erebidae) larvae. Environ Entomol 43(3):595–604.

26. Kieft TL, Simmons KA (2015) Allometry of animal-microbe interactions and global census of animal-associated microbes. Proc R Soc B 282:20150702.

27. Nechvatal JM, et al. (2008) Fecal collection, ambient preservation, and DNA extraction for PCR amplification of bacterial and human markers from human feces. J Microbiol Methods 72(2):124–132.

